# Kidney injury molecule-1 is a potential receptor for SARS-CoV-2

**DOI:** 10.1101/2020.10.09.334052

**Authors:** Chen Yang, Yu Zhang, Hong Chen, Yuchen Chen, Dong Yang, Ziwei Shen, Xiaomu Wang, Xinran Liu, Mingrui Xiong, Kun Huang

## Abstract

COVID-19 patients present high incidence of kidney abnormalities, which are associated with poor prognosis and high mortality. Identification of SARS-CoV-2 in kidney of COVID-19 patients suggests renal tropism and direct infection. Presently, it is generally recognized that SARS-CoV-2 initiates invasion through binding of receptor-binding domain (RBD) of spike protein to host cell-membrane receptor ACE2, however, whether there is additional target of SARS-CoV-2 in kidney remains unclear. Kidney injury molecule-1 (KIM1) is a transmembrane protein that drastically up-regulated after renal injury. Here, binding between SARS-CoV2-RBD and the extracellular Ig V domain of KIM1 was identified by molecular simulations and co-immunoprecipitation, which was comparable in affinity to that of ACE2 to SARS-CoV-2. Moreover, KIM1 facilitated cell entry of SARS-CoV2-RBD, which was potently blockaded by a rationally designed KIM1-derived polypeptide. Together, the findings suggest KIM1 may mediate and exacerbate SARS-CoV-2 infection in a ‘vicious cycle’, and KIM1 could be further explored as a therapeutic target.

## 1. Introduction

The World Health Organization has announced the coronavirus disease 2019 (COVID-19) as a pandemic^1^. Severe acute respiratory syndrome coronavirus 2 (SARS-CoV-2), the pathogen of COVID-19, is a single-stranded RNA virus, belonging to the beta-coronavirus genus which also includes severe acute respiratory syndrome coronavirus (SARS-CoV) and middle east respiratory syndrome coronavirus (MERS-CoV)^2^ SARS-CoV-2, SARS-CoV and MERS-CoV mainly target respiratory systems to primarily manifest with respiratory illness. Notably, reports about renal involvement among patients infected, as well as identifications of viral infection in kidney suggested that these coronaviruses may directly invade kidney^3 4 5^.

Kidney impairment in hospitalized COVID-19 patients is common, and is associated with severe inflammation, poor clinical progress and high in-hospital mortality. A study from Wuhan showed that 251 (75.4%) of 333 COVID-19 patients had renal complications including proteinuria, hematuria or acute kidney injury (AKI), and patients with renal impairment had much higher overall mortality (11.2%) than those without (1.2%)^6^. Meanwhile, in a cohort of 5449 patients from New York, 36.6% of hospitalized patients developed AKI, and 35% of those with AKI died during observation time^7^. Consistently, we recently reported that COVID-19 patients with chronic kidney diseases (CKD) were related to higher risk of poor prognosis and in-hospital death^8, 9^. A recent study suggested renal tropism of SARS-CoV-2, which was detected in the kidneys of 72% of COVID-19 patients with AKI, whereas the viral RNA was only found in 43% of patients without AKI^5^. Among multiple organ manifestations in COVID-19 patients, apart from lung, kidney is highly vulnerable to the virus, and renal dysfunctions are closely associated with high mortality, with the underlying molecular mechanisms remain unclear.

Like coronaviruses SARS-CoV and MERS-CoV, SARS-CoV-2 contains four key proteins, namely spike (S), envelope (E), membrane (M) and nucleocapsid (N) proteins. SARS-CoV-2 invasion initiates from binding with cellular membrane receptors via the viral spike protein^2^ (Figure 1A). Presently, angiotensin-converting enzyme 2 (ACE2), also the target for SARS-CoV, is the only confirmed receptor for SARS-CoV-2^2^. Responsible for receptor recognition, SARS-CoV-2 spike protein (SARS-CoV-2-SP) consists subunits S1 and S2, the receptor-binding domain (RBD) in S1 binds ACE2 to initiate the fusion of S2 with cell membrane and subsequent cell entry^10^ (Figure 1A). Renal expression of ACE2 has been identified^10^, suggesting its potential of mediating SARS-CoV-2 kidney infection. Cell entry of virus may involve multiple transmembrane receptors^11^, whether there are any additional receptors in kidney remain unclear.

**Figure 1.**
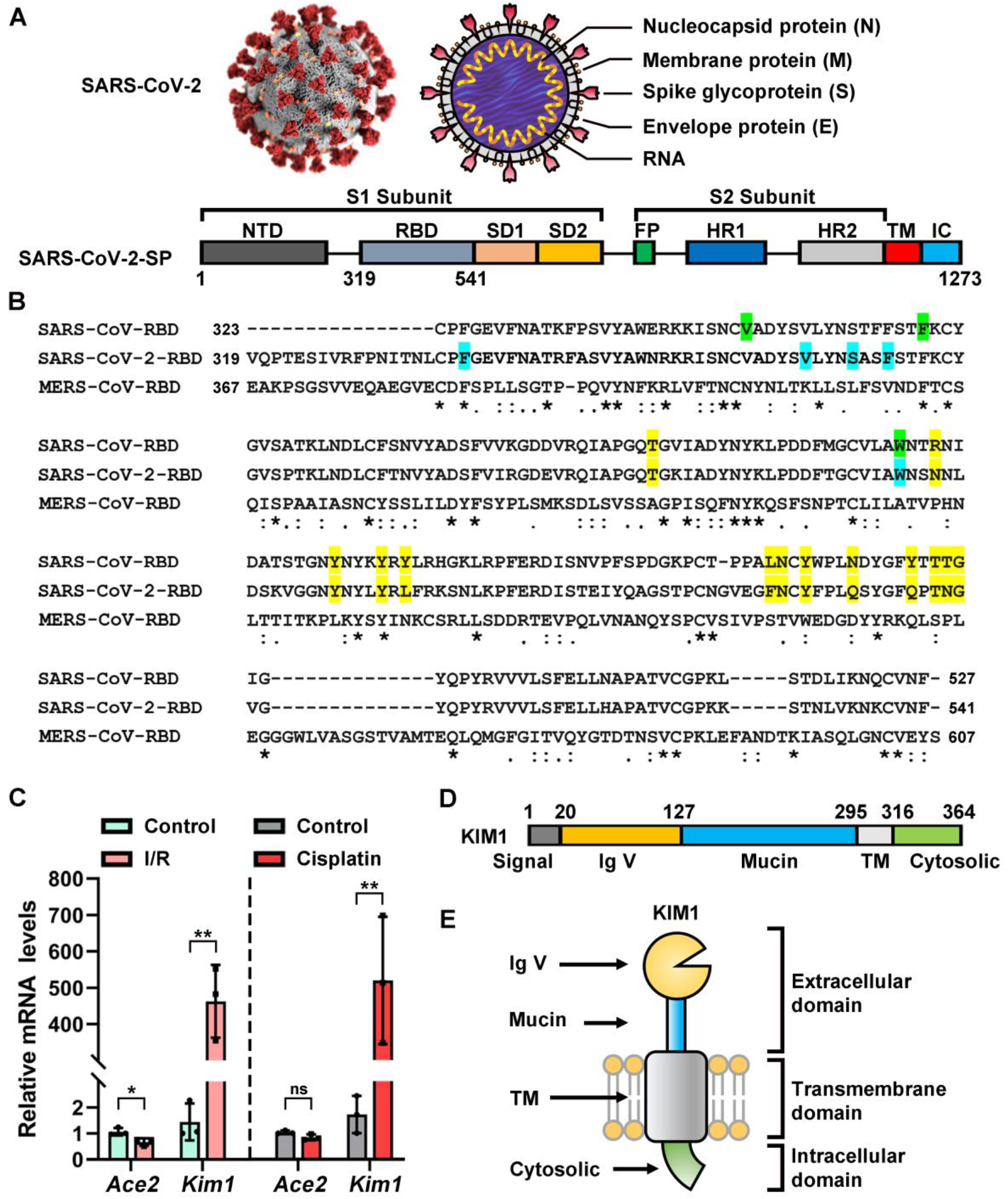
Basic information of SARS-CoV-2-SP and KIM1. (A) Structural scheme of SARS-CoV-2 spike protein. NTD, N-terminal domain; RBD, receptor-binding domain; RBM, receptor-binding motif; SD1, subdomain 1; SD2, subdomain 2; FP, fusion peptide; HR1, heptad repeat 1; HR2, heptad repeat 2; TM, transmembrane region; IC, intracellular domain. (B) Sequence alignment of the receptor-binding domain of SARS-CoV, SARS-CoV-2 and MERS-CoV ACE2-contacting residues of SARS-CoV-RBD and SARS-CoV-2-RBD are highlight in yellow; KIM1-contacting residues of SARS-CoV-RBD are in green; KIM1-contcacting residues of SARS-CoV-RBD-2 are in blue; asterisks indicate fully conserved residues; colons indicate partly-conserved residues; periods indicate weakly-conserved residues. (C) Relative mRNA level of ACE2 and KIM1 from the kidneys of ischemia-reperfusion (I/R) and cisplatin induced kidney injury mouse models. \**P* < 0.05, \*\**P* < 0.01; ns, no significance. (D) Structure scheme of KIM1 domains, in relation to cell membrane (E). Signal, signal peptide region; Ig V, immunoglobulin variable Ig-like domain; Mucin, mucin containing domain; TM, transmembrane region.

Kidney injury molecule-1 (KIM1, also known as TIM1, HAVCR1, or CD365) is primary expressed in kidney and drastically up-regulated in injured kidney proximal tubule in AKI or CKD, and plays crucial roles in inflammation infiltration and immune response^12, 13^. Structurally, KIM1 is composed of immunoglobulin variable Ig-like (Ig V) domain, mucin domain, transmembrane domain and cytosolic domain. Among them, Ig V domain mediates Ebola and Dengue invasion by direct binding virus^14, 15^, or as a viral envelope phosphatidylserine (PS) binding receptor to mediate viral invasions that cause Hepatitis A and Encephalitis^16, 17^. Here, we hypothesize that KIM1 may be a binding target of SARS-CoV-2 that mediates its kidney invasion.

In this study, we investigate the binding of SARS-CoV2-RBD to KIM1 by molecular simulations and co-immunoprecipitation (Co-IP) assays, binding potentials of KIM1 to SARS-CoV and MERS-CoV were also evaluated. We demonstrate KIM1-assisted cell entry and cytotoxicity of SARS-CoV2-RBD in human kidney cells, and explore the blockade effects of two KIM1-derived antagonist peptides against SARS-CoV-2-RBD invasion.

## 2. Materials and Methods

### 2.1 Materials

Recombinant SARS-CoV-2-RBD (T80302) was obtained from Genscript (NanJing, China). Antagonist peptide 1 (SCSLFTCQNGIV) and 2 (SCSLFTCQNGGGWF) were chemically synthesized by Genscript. Anti-Mouse-IgG (H&L) antibody (p/n 18-8817-33) and anti-Rabbit-IgG (H&L) antibody (p/n 18-8817-33) were obtained from RockLand (Philadelphia, PA). IgG with SureBeads™ Protein G magnetic beads (J2112LB-02) were purchased from Bio-Rad (Hercules, CA). DAPI (D9542) was from Sigma (St. Louis, MO). Alex Flour 594 labelled phalloidin (C2205S) was from Beyotimes (Shanghai, China). Antibodies against KIM1 (NBP1-76701, Novus Biologicals, New York, NY), Flag (F1804, Sigma), HA (H6908, Sigma), ACE2 (21115-1-AP, Proteintech, Wuhan, China) were used.

### 2.2 Acquisition and analysis of transcriptome sequencing data of KIM1 and ACE2

To obtain the comprehensive transcriptome and protein profiles of KIM1 and ACE2 for human tissues, we collected and analyzed the transcriptome data and immunohistochemistry-base protein profiles from Human Protein Atlas (HPA, https://www.proteinatlas.org), which showed the expression and localization of human proteins across tissues and organs, based on deep sequencing of RNA (RNA-seq) from 37 normal tissue and immunohistochemistry on tissue microarrays containing 44 tissue types^18^. HPA RNA-seq tissue of the protein-coding gene was recorded as mean protein-coding transcripts per million (pTPM), corresponding to mean values of samples from each tissue. Histology-based protein expression levels were analyzed manually into 4 levels (not detected, low, medium and high). Top 10 tissular transcriptional levels and histology-based protein expression levels of KIM1 and ACE2 were listed, respectively. And the overlapped expression profile of KIM1 and ACE2 were summarized.

### 2.3 Molecular docking and dynamics simulations

Dockings were conducted *via* Z-Dock (http://zdock.umassmed.edu/). Crystal structures of SARS-CoV-RBD (PBD ID 2AJF), SARS-CoV-2-RBD (PDB ID 6M0J), MERS-COV-RBD (PDB ID 4L3N), ACE2 (PDB ID 1R42) and KIM1 Ig V (PDB ID 5DZO) were used to seek potential binding models. The best-scored protein complexes were selected to conduct following molecular dynamics simulations, which were conducted by the Desmond server (http://www.deshawresearch.com/) and analyzed by Pymol2.3 and Maestro11.8.012. 50 nsec dynamics simulations diagram was applied to study the dynamic parameters of the protein complexes. Molecular Mechanics Generalized Born Surface Area (MM-GBSA) binding free energy was calculated by HawkDock^19^.

### 2.4 Root Mean Square Deviation and Root Mean Square Fluctuation

Root Mean Square Deviation (RMSD) was utilized to estimate the average change in displacement of a selection of atoms for a particular frame as described^20^. Root Mean Square Fluctuation (RMSF) was conducted to study the displacement changes in the protein chain^20^.

### 2.5 AKI mouse models and qPCR

I/R injury was performed on C57BL/6 mice as we previously described^21^. For cisplatin-induced AKI, 30 mg/kg bodyweight cisplatin was injected intraperitoneally into 8-week-old male mice, and mice were sacrificed 3 days later. Blood and kidney samples were collected for further analysis. Total RNA was isolated from kidneys by RNA^iso^ Plus (TaKaRa Biotech., Dalian, China) and reverse transcribed into cDNA using the M-MLV first-strand synthesis system (Invitrogen, Grand Island, NE). The abundance of specific gene transcripts was assessed by qPCR. Primers used in the study are provided (Supplementary Table S4).

### 2.6 Constructs

Mammalian expression plasmids for human KIM1, KIM1 Ig V (KIM1 20-127 aa), KIM1 ΔIg V (truncated KIM1 without residues 20-127 aa), ACE2, SARS-CoV-2-RBD (SARS-CoV-2 spike protein 319-541 aa) and SARS-CoV-2-SP were constructed. PCR amplification products of the corresponding cDNA fragments were cloned into a pRK promoter-based vector containing either HA or FLAG tag (Figure 3A). SARS-CoV-2 related plasmids were kindly gifts by Dr. PH Wang of Shandong University.

### 2.7 Cell culture and transfection

Human kidney tubular cell line HK-2 (obtained from China Center for Type Culture Collection, Wuhan, China) was cultured in DMEM/F12 media (Hyclone, Logan, UT) containing 17.5 mM glucose and 10% Fetal Bovine Serum. To evaluate the impact of SARS-CoV-2 on cells, HK-2 cells were transfected with SARS-CoV-2-SP and SARS-CoV-2-RBD plasmids for 24 hours, then collected for further detection.

### 2.8 Co-immunoprecipitation (Co-IP)

Transfected HK-2/HEK293T cells were lysed in 1 mL pre-lysis buffer. For immunoprecipitation, cell lysate was immune-precipitated with indicated antibody or respective IgG with SureBeads™ Protein G Magnetic Beads overnight at 4 °C. After washing with pre-lysis buffer containing 500 mM NaCl, the beads were boiled in loading buffer and subjected to immunoblotting.

### 2.9 CRISPR-Cas9-mediated knockout of KIM1

The CRISPR-Cas9 based protocols for genome engineering were used as we previously described^22^. KIM1 guide RNA target sequences are provided (Table S4).

### 2.10 Fluoresceine isothiocyanate label

Fluoresceine isothiocyanate (FITC) label was performed as we previously described^23^. Briefly, SARS-CoV-2-RBD was co-incubated with FITC (molar ratio 1:5) overnight, then 5 mM NH_4_Cl was added to stop the reaction and quench the un-reacted FITC. The solution was dialyzed twice, and lyophilized for further use.

### 2.11 Confocal microscopy

Transfected HEK293T cells (5 × 10^6^) or HK-2 cells (1 × 10^7^) were incubated with free FITC or FITC-SARS-CoV-2-RBD for 2 h. For peptide based internalization assays, antagonist peptide 1 or 2 (50 μM) was co-added with SARS-CoV-2-RBD (100 μg/mL). After fixing with 4% (W/V) formaldehyde, the cell membranes were stained with Alex Flour 594 labelled phalloidin (2 μg/mL) and the nuclei were stained by DAPI (1 μg/mL), then imaged with a Leica TCS SP8 confocal microscope.

### 2.12 Cell Viability Assays

Cells were plated at 3000-4000 cells per well. At 80% confluence, cells incubated with SARS-CoV-2-RBD (100 μg/ml) were treated with or without antagonist peptide 1/2 (50 μM). After that, 10 μL MTT (5 mg/ml) was added to each well for 4 hours, medium was removed and DMSO was added. Absorbance measured at 490 nm was normalized to the respective control group.

### 2.13 Statistical analysis

Data were expressed as mean ± SD. Significant differences were assessed by two-tailed Student’s test. 2-sided P value less than 0.05 was considered statistically significant. Analyses were performed with EXCEL 2017 and GraphPad Prism 8.0.

## 3. Results

### 3.1 Expression profiles of KIM1 and ACE2 in human tissues

To elucidate KIM1 and ACE2 enrichment in tissues, the transcriptome and histology-based protein expression data from the Tissue Atlas of Human Protein Atlas were collected. Top 10 tissues with mRNA and protein abundance of KIM1 and ACE are listed (Figure S1A-F). Notably, KIM1 and ACE2 co-expressed in kidney, colon, rectum, testis and gallbladder (Figure S1E-F), which are all among the target organs of SARS-CoV-2^24, 25^, implicating a potential close correlation of KIM1 with COVID-19 manifestations.

### 3.2 Molecular dockings reveal interaction between SARS-CoV-2 RBD and KIM1 Ig V

The overall sequence similarity between SARS-CoV-RBD and SARS-CoV-2-RBD is 62.93%, and 9 of the 14 ACE2-contacting residues are conserved in both RBD (Figure 1B), indicating possible invasion via the identical receptor. In comparison, MERS-CoV-RBD, which recognizes DPP4^26^, shows low sequence similarity with SARS-CoV-RBD and SARS-CoV-2-RBD (17.07% and 14.86%, respectively; Figure 1B).

Consistent with a previous report^27^, *Kim1* is drastically up-regulated in the kidneys of ischemia-reperfusion- or cisplatin-injured mice, while only mild changes of *Ace2* were observed under the same stress (Figure 1C). Among four domains of KIM1 (Figure 1D-E), Ig V domain is responsible for virus binding^14, 15^, molecular dynamic docking was thus conducted to investigate its binding with SARS-CoV-2-RBD.

We evaluated the equilibrium of docking diagram to ensure reliable calculations. RMSD and RMSF demonstrated minor displacement, indicating the docking is equilibrated (Figure S1G-I). Structural information of SARS-CoV-2-RBD and KIM1 Ig V complex was provided in Figs. S2A-S2C. MM-GBSA analysis suggested that Phe338, Val367, Ser371, Phe374 and Trp436 of SARS-CoV-2-RBD interact with Leu54, Phe55, Gln58, Trp112 and Phe113 of KIM1 Ig V (Figure 2A). Moreover, Phe374 and Trp436 of SARS-CoV-2-RBD bind with Phe55 of KIM1 Ig V to form π-π conjugation, while Val367 of SARS-CoV-2-RBD bind with Phe113 and Leu54 of KIM1 Ig V via hydrophobic interaction, indicating close contacts between SARS-CoV-2-RBD and KIM1 (Figure 2A). Above interactions contribute to a combined binding free energy of −35.64 kcal/mol (Table S1 and S2), which is lower than that of SARS-CoV-2-RBD and ACE2 (−50.60 kcal/mol), but comparable to that of SARS-CoV-RBD and ACE2 (−38.3 kcal/mol)^26^. Given ACE2 is also recognized as a key receptor for SARS-CoV-RBD^26^, our results thus indicate strong interaction between SARS-CoV-2-RBD and KIM1 (Table 1 and Table S1). Notably, the different binding regions on the surface of SARS-CoV-2-RBD to KIM1 and to ACE2 implicate that KIM1 and ACE2 may synergistically mediate SARS-CoV-2 invasion (Figure 2B).

**Figure 2.**
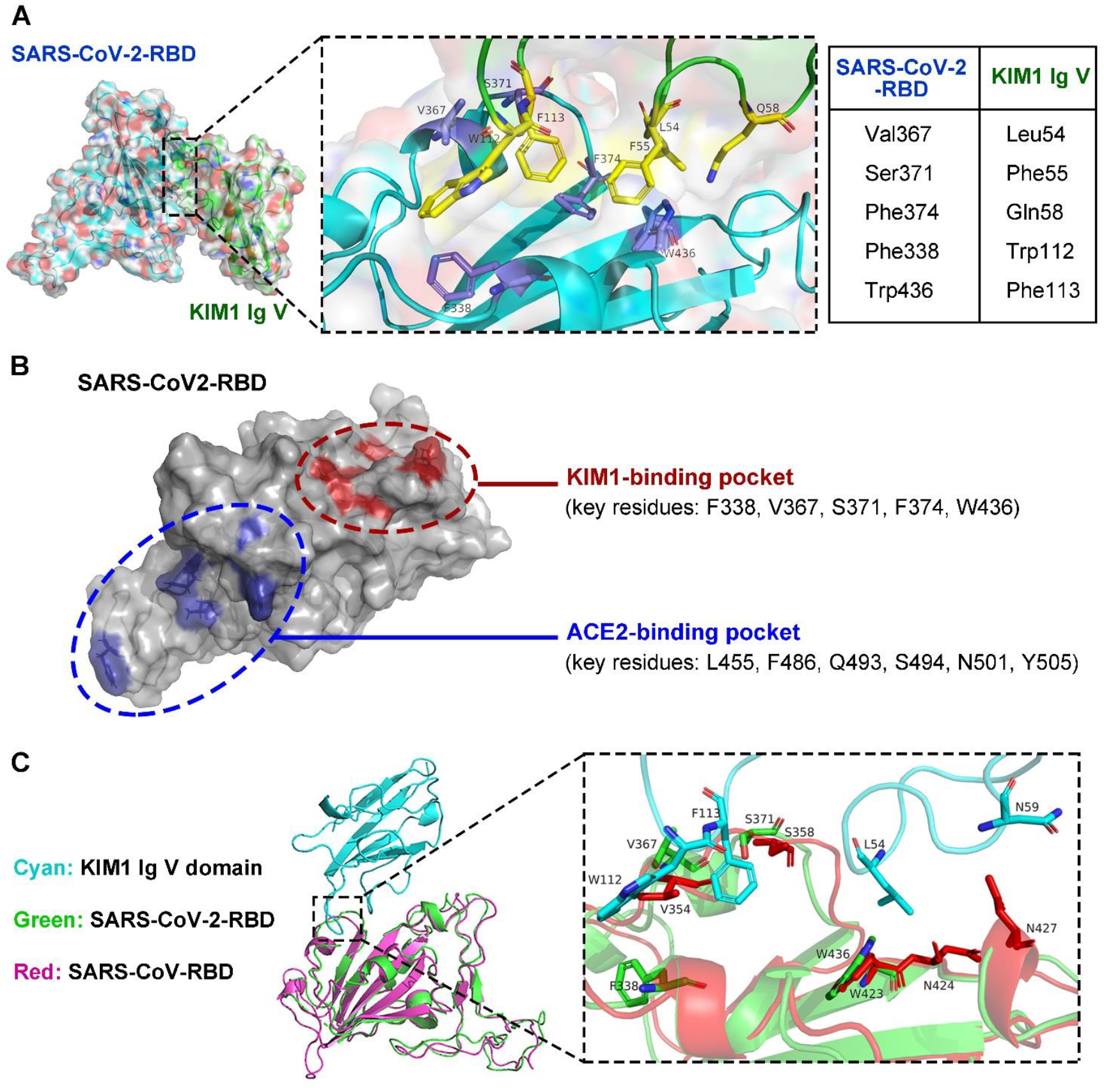
Binding model of SARS-CoV-2-RBD and KIM1 Ig V. (A) Low-energy binding conformations of SARS-CoV-2-RBD binds to KIM1 Ig V. Left panel, the surface model of SARS-CoV-2-RBD. Right panel, the high-resolution image of the binding sites, Phe338, Val367, Ser371, Phe374, and Trp436 of SARS-CoV-2-RBD interact with Leu54, Phe55, Gln58, Trp112 and Phe113 of KIM1 Ig V. (B) Distinct binding regions of KIM1 and ACE2 in SARS-CoV-2-RBD, with KIM1-binding pocket in red and ACE-2 binding pocket in blue. (C) SARS-COV-RBD and SARS-CoV-2-RBD bind with the same pocket of KIM1 Ig V.

**Table 1.** MM-GBSA binding free energy of residues in SARS-CoV-2-RBD and KIM1 Ig V complex.

| Rank | SARS-CoV-2-RBD |  | KIM1 Ig V |  |
| --- | --- | --- | --- | --- |
|  | Residue | Binding Free Energy<br>(kcal/mol) | Residue | Binding Free Energy<br>(kcal/mol) |
| 1 | Val367 | -2.42 | Leu54 | -6.14 |
| 2 | Trp436 | -2.00 | Trp112 | -5.26 |
| 3 | Phe374 | -1.86 | Leu54 | -4.96 |
| 4 | Ser371 | -1.84 | Phe113 | -3.97 |
| 5 | Phe338 | -1.53 | Gln58 | -2.11 |
| 6 | Leu368 | -1.50 | Asn114 | -0.75 |
| 7 | Asn437 | -1.46 | Asn59 | -0.13 |
| 8 | Asn440 | -1.46 | Asp115 | -0.13 |
| 9 | Ser373 | -1.18 | Asp74 | -0.12 |
| 10 | Leu335 | -0.96 | Asp99 | -0.11 |

Top 10 ranked residues that involved in the binding of SARS-CoV-2-RBD and KIM1 Ig V were listed.

Global population sequencing data have revealed a large number of mutations in SARS-CoV-2 spike protein (Figure S3 and Table S3), which may affect the interaction between SARS-CoV-2 and its receptors. COVID-19 cases carrying V367F mutation in SARS-CoV-2, which contacting KIM1, have been reported (http://giorgilab.dyndns.org/coronapp/, Figure S3B-C). MM-GBSA analysis suggest that V367F mutation lead to mildly enhanced binding free energy (−37.26 kcal/mol) to KIM1 compared with that of SARS-CoV-2-RBD (−35.64 kcal/mol) (Table S1). Consistent with our findings, latest clinical reports combined with mutational scanning suggest that V367F may lead to enhanced viral infectivity of SARS-CoV-2^28, 29^.

### 3.3 SARS-CoV-RBD and SARS-CoV-2-RBD target the same binding pocket in KIM1

Microarray data show increased KIM1 expression in SARS patients-derived peripheral blood mononuclear cells comparing to healthy controls (Figure S4A) (GSE1739)^30^. Considering SARS-CoV-RBD and SARS-CoV-2-RBD were both reported to invade kidney^3, 31^, and share high homology (Figure 1B), we further evaluated the binding potential of SARS-CoV-RBD with KIM1 (Figure S4-S5). SARS-CoV-RBD binds KIM1 Ig V at a combined binding free energy of −21.59 kcal/mol (Tables S1 and S2), suggesting a relatively lower affinity to KIM1 than that of SARS-CoV-2-RBD (−35.64 kcal/mol). Val354, Phe360, Trp423 and Asn424 of SARS-CoV-RBD interact with Leu54, Phe55, Gln58, Trp112 and Phe113 of KIM1 Ig V (Figure S5C), suggesting that both coronaviruses share the same binding pocket within KIM1. In addition, MM-GBSA binding free energy between MERS-COV-RBD and KIM1 Ig V is −10.12 kcal/mol, indicating a weak interaction (Table S1). Together, our data suggest that compared with SARS-CoV-RBD and MERS-COV-RBD, SARS-CoV-2-RBD showed the highest binding affinity to KIM1; moreover, SARS-CoV-RBD and SARS-CoV-2-RBD share the same binding pocket on the KIM1 Ig V domain (Figure 2C).

### 3.4 Intracellular interaction of SARS-CoV-2-RBD and KIM1 Ig V

To confirm the binding between SARS-CoV-2-RBD and KIM1, endogenous and exogenous Co-IP assays were performed (Figure 3A). To mimic renal cells under exposure of SARS-CoV-2, a human kidney tubular epithelial cell line HK-2 was transfected with plasmid expressing SARS-CoV-2-SP or SARS-CoV-2-RBD; the binding between KIM1 and SARS-CoV-2-SP or SARS-CoV-2-RBD were confirmed (Figure 3B). Similar results were observed in exogenous Co-IP assays in HEK293T, and comparable binding of SARS-CoV-2-RBD to KIM1 and to ACE2 were suggested (Figure 3C).

**Figure 3.**
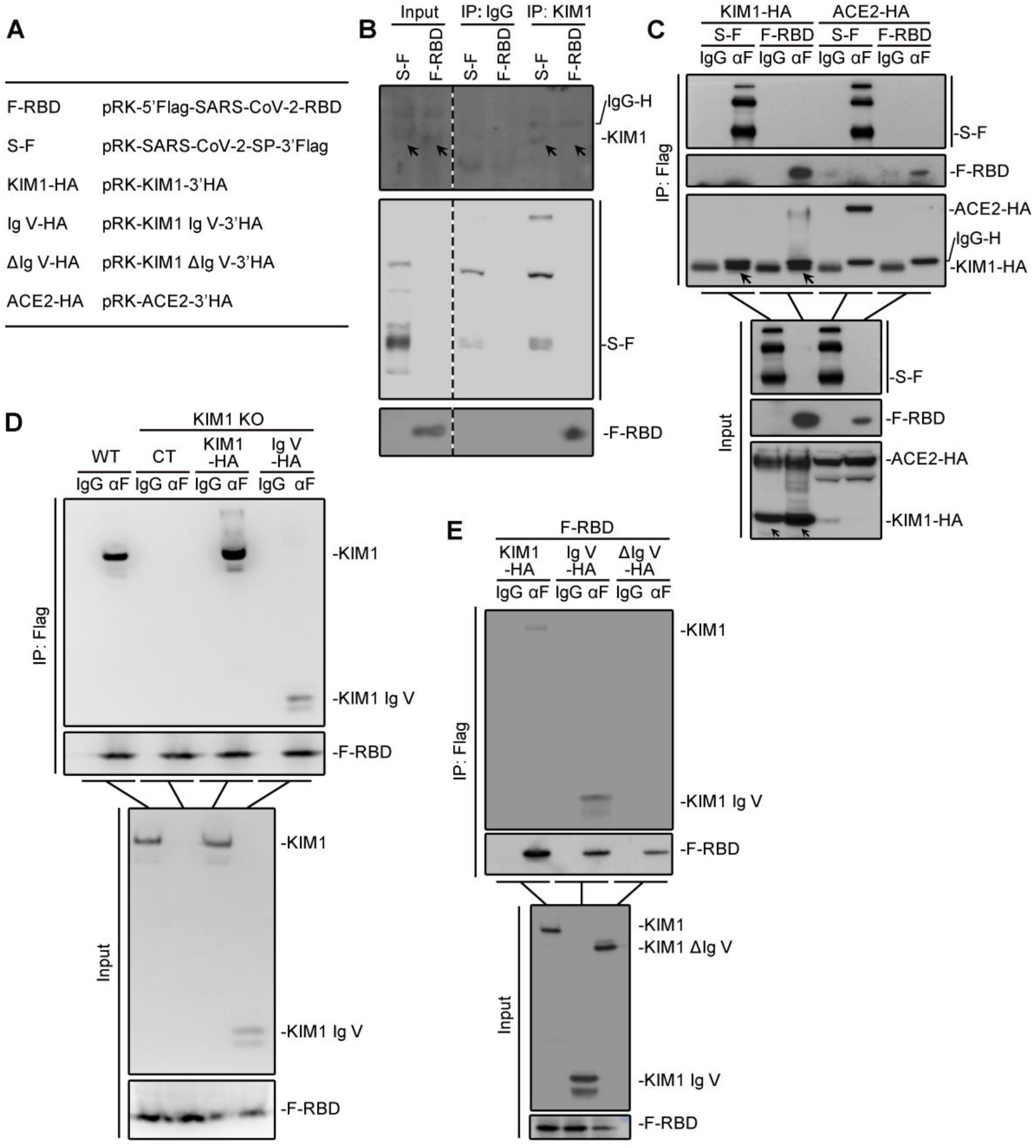
SARS-CoV-2-RBD binds with KIM1 Ig V. (A) Constructs used in Co-IP studies. (B) Endogenous Co-IP results in HK-2 cell line. (C) Exogenous Co-IP results in HEK393T. (D) Detection of the binding between KIM1 Ig V and SARS-CoV-2-RBD in KIM1 knockout HK-2 cell line. For IP group, KIM1 and KIM1 Ig V was detected by anti-KIM1 antibody. (E) Detection of the binding between KIM1 Ig V and SARS-CoV-2-RBD in HEK293T. For IP group, KIM1 (HA-tag), KIM1 ΔIg V (HA-tag) and KIM1 Ig V (HA-tag) were detected by anti-HA antibody.

Since KIM Ig V is crucial in mediating the binding of virus and cell membrane receptor^14 32^, full length KIM1 and Ig V were respectively co-transfected with SARS-CoV-2-RBD into a stable KIM1 knockout HK-2 cell line (Figure 3D and Figure S6A-B); binding of SARS-CoV-2-RBD with full length KIM1 and Ig V were observed (Figure 3D). Deletion of Ig V domain abolished the binding between KIM1 and SARS-CoV-2-RBD in HEK293T cells (Figure 3E). These findings suggest Ig V domain mediates interaction between KIM1 and SARS-CoV-2.

### 3.5 KIM1 mediates the cell entry of SARS-CoV-2-RBD

We next investigated the role KIM1 plays in mediating cell entry of SARS-CoV-2-RBD. Fluoresceine isothiocyanate (FITC) was utilized to label SARS-CoV-2-RBD, and the cell entry of SARS-CoV-2-RBD in HK-2 cells was observed (Figure 4A). Knockout of KIM1 attenuated the invasion and reduced the cytotoxicity induced by SARS-CoV-2-RBD (Figure 4A and Figure S6C). In KIM1 knockout HK-2 cells, restoring full length KIM1 and overexpressing Ig V both rescued cell entry of SARS-CoV-2-RBD (Figure 4A). Consistently, overexpression of KIM1 in HEK293T leads to enhanced invasion of SARS-CoV-2-RBD (Figure 4B).

**Figure 4.**
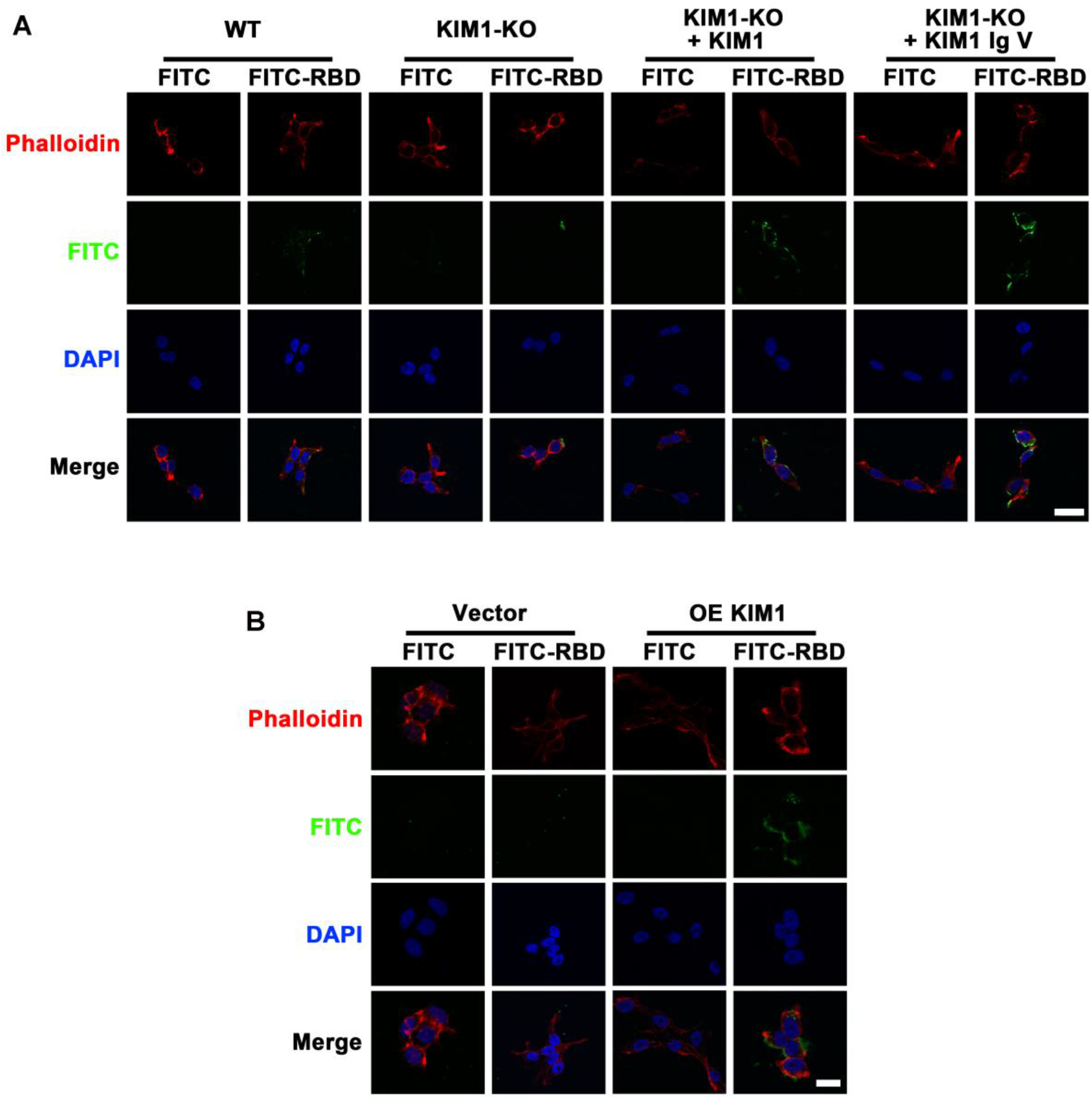
KIM1 mediates the cell entry of SARS-CoV-2-RBD. (A) Cell entry of SARS-CoV-2-RBD in WT and KIM1 knockout HK-2 cell line. (B) Cell entry of SARS-CoV-2-RBD in HEK293T. Scale bar, 20 μm.

### 3.6 A KIM1-derived peptide inhibits the cell entry of SARS-CoV-2-RBD

To competitively bind with SARS-CoV-2-RBD and inhibit KIM1-associated invasion, we rationally designed two antagonist peptides based on SARS-CoV-2 contacting motifs in KIM1 (motif 1: Leu54, Phe55, Gln58; motif 2: Trp112, Phe113; Fig 5A). Peptide 1 mimics motif 1, while peptide 2 covers both motifs, with triple glycine used as a flexible linker (Figure 5A). The MM-GBSA binding free energy which indicates potential binding between peptides and SARS-CoV-2-RBD was provided in Table S1. Both peptides showed no distinct cytotoxicity, and peptide 2 reduced SARS-CoV-2-RBD induced cell death (Figure S6D) and decreased cell entry of SARS-CoV-2-RBD in HK-2 cells (Figure 5B), suggesting the protective effects of this KIM1-derived polypeptide.

**Figure 5.**
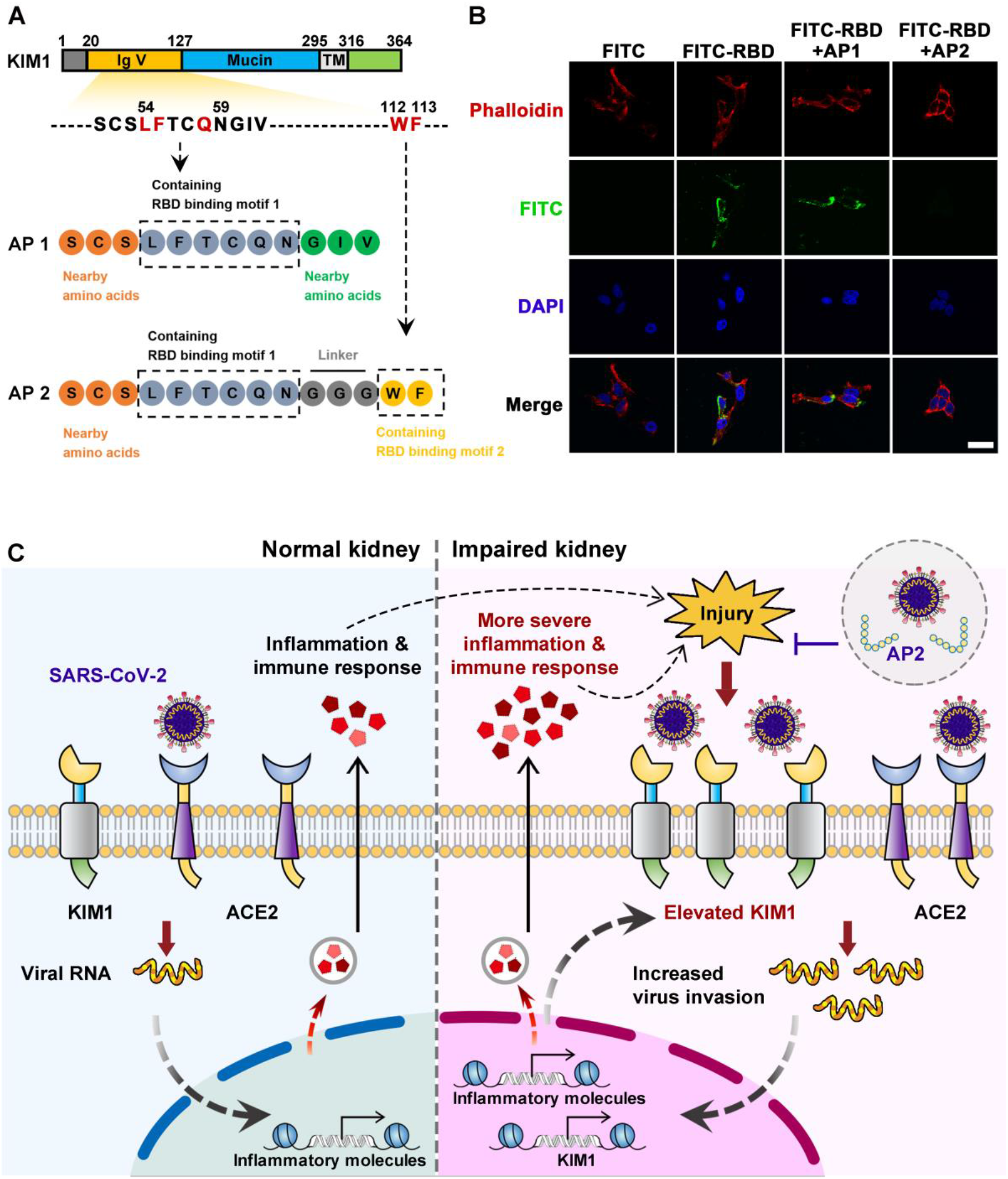
Rationally designed antagonist peptide 2 inhibits the cell entry of SARS-CoV-2. (A) Schematic diagram of antagonist peptides 1 and 2. (B) Effects of antagonist peptide 1 (AP1) and antagonist peptide 2 (AP2) on SARS-CoV-2-RBD entry. Scale bar, 20 μm. (C) A vicious cycle in the kidney of COVID-19 patients mediated by KIM1. KIM1/ACE2 mediates the entry of SARS-CoV-2 and leads to the initial direct kidney infection, the resultign kidney injury drastically up-regulates KIM1, thus in turn promotes the secondary infection and consequently exacerbating the kidney damage. KIM1-derived antigonist peptide (AP) may competitively bind with SARS-CoV-2-RBD to hinder viral invasion.

## 4. Discussion

To fight COVID-19 pandemic, a deep understanding of how SARS-CoV-2 invade human cells is warranted. Studies have indicated direct infection of SARS-CoV-2 in kidney in addition to lung^5, 31^, however, ACE2 remains the only confirmed receptor which may mediate this invasion. Furthermore, the renal tropism of SARS-CoV-2 and associated kidney injury, seem unexplainable by the relatively decreased level of ACE2 upon viral invasion^33^. Here, we reported KIM1, a drastically up-regulated biomarker for kidney injury (Figure 1C)^34^, mediates SARS-CoV-2 kidney invasion as a receptor. We also found that SARS-CoV-2-RBD binds KIM1 Ig V with a higher affinity than that of SARS-CoV-RBD and MERS-COV-RBD, which probably underlies the stronger contagion of SARS-CoV-2^35^. Notably, our results suggest that SARS-CoV-2-RBD binds KIM1 and ACE2 *via* two distinct pockets, implicating that KIM1 and ACE2 may synergistically mediate the invasion of SARS-CoV-2 in kidney cells; which may explain the strong renal tropism, as well as the high incidence of acute kidney injury in COVID-19 patients^5^.

Mutations in SARS-CoV-2 spike protein may affect the interaction between SARS-CoV-2 and its receptors, resulting in different viral infectivity and antigenicity^28^. Among the sites of SARS-CoV-2 that involved in attachment to KIM1, V367F mutation was recently reported to increase sensitivity to neutralizing antibodies and show stability-enhancing effect^28, 29^. Our findings suggested that V367F mutation enhances binding affinity towards KIM1 (Table S1), yet the functional consequences remain unknown; further investigations of how KIM1 interacts with different SARS-CoV-2 variants could be of value in developing related therapies.

As a viral envelope PS binding receptor, KIM1 has been shown to mediate viral invasions that cause Hepatitis A and Encephalitis through its PS binding site WFND (112-115)^15, 16, 32^. Although our results have demonstrated the binding of KIM1 with RBD of multiple coronaviruses, its role as a PS receptor in meditating SARS-CoV-2 binding also worth further investigation.

ACE2 is the most well-studied receptor for SARS-CoV-2 so far, yet it is not an ideal therapeutic target for COVID-19 since it is widely expresses in multiple organs, and plays crucial roles in regulating blood pressure and preventing heart/kidney injury^36, 37^. In contrast, KIM1 has stronger association to kidney function and highly expressed only after renal injury^14, 16, 17^, which make it a more specific and maybe safer therapeutic target for COVID-19 patients with kidney diseases.

Together, our study identified KIM1, a kidney injury marker, as a potential receptor for SARS-CoV-2. To explain the high incidence of AKI of COVID-19 patients, we propose a model of a ‘vicious cycle’ co-mediated by KIM1 and ACE2 in kidney of COVID-19 patients. In this model, the higher physiological level and binding affinity of ACE2 make it the primary target for the initial SARS-CoV-2 invasion, which lacks kidney-specificity. Next, the induced kidney injury and the resulting drastically up-regulated KIM1 rapidly promotes a KIM1-and-ACE2-co-mediated secondary viral infection, which is more kidney-specific, and consequently exacerbating kidney damage in a vicious cycle (Figure 5C). Therefore, blocking KIM1 may partly inhibit the cell entry of SARS-CoV-2 and attenuate the kidney injury caused by viral invasion, however, further studies are necessary to fully validate this model. Moreover, drugs target the binding pocket of KIM1 may be explored, and bi-specific antibodies/peptides that dual-targeting KIM1 and ACE2 may also be developed, considering that they can be simultaneously targeted by SARS-CoV-2. Given expression profile of KIM1 overlapped with that of ACE2, notably in multiple target organs of SARS-CoV-2 (Figure S1), studies on additional organs/tissues affected by coronaviruses may also worth future exploration.

## Abbreviations

AKI: Acute kidney injury
CKD: Chronic kidney diseases
KIM1: Kidney injury molecule-1
Co-IP: Co-immunoprecipitation
HPA: Human Protein Atlas
MERS-CoV: Middle east respiratory syndrome coronavirus
RBD: Receptor-binding domain
RMSD: Root Mean Square Deviation
RMSF: Root Mean Square Fluctuation
SARS-CoV: Severe acute respiratory syndrome coronavirus
SARS-CoV-2: Severe acute respiratory syndrome coronavirus 2
SARS-CoV-2-SP: Severe acute respiratory syndrome coronavirus spike protein

## Acknowledgements

This work was supported by the Natural Science Foundation of China (31671195 and 31971066).

## Author contributions

Chen Yang Yu Zhang, Hong Chen and Dong Yang performed the simulation studies. Chen Yang, Yu Zhang, Hong Chen and Yuchen Chen performed the Co-IP. Chen Yang, Yu Zhang, Hong Chen, Mingrui Xiong and Xinran Liu performed the confocal assays. Chen Yang, Yu Zhang, Hong Chen, Ziwei Shen and Xiaomu Wang analyzed the data. Chen Yang, Yu Zhang, Hong Chen and Yuchen Chen draft the manuscript. Kun Huang designed the study and revised the manuscript. All authors have approved the manuscript.

## Data availability

All other data are available from the corresponding authors on reasonable request.

