## Supplemental materials for "Kidney injury molecule-1 is a potential receptor for SARS-CoV-2"

*Short Title: KIM1 mediates SARS-CoV-2 invasion*

Chen Yang[1#], Yu Zhang[1#], Hong Chen[1#], Yuchen Chen[1], Dong Yang[1], Ziwei Shen[1], Xiaomu Wang[1], Xinran Liu[1], Mingrui Xiong[1] & Kun Huang[1,2*]

[1]School of Pharmacy, Tongji Medical College, Huazhong University of Science & Technology, Wuhan, China, 430030

[2]Tongji-RongCheng Biomedical Center, Tongji Medical College, Huazhong University of Science & Technology, Wuhan, China, 430030

[#] Equal contribution

*Corresponding author

Kun Huang, Ph.D.

Tongji School of Pharmacy

Tongji Medical College

Huazhong University of Science & Technology

Wuhan, China, 430030

### Supplemental materials Table of Contents

Table S1. MM-GBSA binding free energy of SARS-CoV-2-RBD and receptors.

Table S2. MM-GBSA binding free energy of residues in SARS-CoV-RBD and KIM1 Ig V complex.

Table S3. Top 20 clinically identified mutations in SARS-CoV-2 spike protein.

Table S4. Primers used in the study.

Figure S1. Expression profiles of KIM1, ACE2 and molecular dynamics docking information of SARS-CoV-2-RBD and KIM1.

Figure S2. Molecular dynamics simulations information of SARS-CoV-2-RBD and KIM1.

Figure S3. Clinically identified mutations in SARS-CoV-2 spike protein

Figure S4. Molecular dynamics simulations information of SARS-CoV-RBD and KIM1.

Figure S5. Binding model of SARS-CoV-RBD and KIM1.

Figure S6. Identification of KIM1 knockout HK-2 cell line and the protective effects of antagonist peptide on SARS-CoV-2-RBD induced cytotoxicity.

**Table S1.** MM-GBSA binding free energy of SARS-CoV-2-RBD and receptors.

| Ligand | Receptor | Binding Free Energy （kcal/mol） |
|---|---|---|
| SARS-CoV-RBD | KIM1 Ig V domain | -21.59 |
| SARS-CoV-2-RBD | KIM1 Ig V domain | -35.64 |
| SARS-CoV-2-RBD (V367F) | KIM1 Ig V domain | -37.26 |
| SARS-CoV-2-RBD | ACE2 | -50.60 |
| MERS-COV-RBD | KIM1 Ig V domain | -10.12 |
| SARS-CoV-2-RBD | AP1 | -7.13 |
| SARS-CoV-2-RBD | AP2 | -6.65 |

MM-GBSA binding free energy of the protein complex was cauclated by the accumulation of the binding free energy of all involved residues. The binding free energy of SARS-CoV-2-RBD and ACE2 was also cauclated and provided as a reference.

**Table S2.** MM-GBSA binding free energy of residues in SARS-CoV-RBD and KIM1 Ig V domain complex.

| Rank | SARS-CoV-RBD | | KIM1 Ig V | |
|---|---|---|---|---|
| | Residue | Binding Free Energy （kcal/mol） | Residue | Binding Free Energy （kcal/mol） |
| 1 | Phe360 | -3.67 | Trp112 | -4.87 |
| 2 | Val354 | -1.93 | Phe55 | -4.33 |
| 3 | Asn424 | -1.34 | Leu54 | -4.17 |
| 4 | Trp423 | -1.12 | Phe113 | -3.62 |
| 5 | Asn427 | -0.71 | Gln58 | -1.46 |
| 6 | Thr359 | -0.69 | Asn114 | -1.42 |
| 7 | Ser358 | -0.68 | Asn59 | -0.54 |
| 8 | Phe361 | -0.59 | Asp115 | -0.37 |
| 9 | Leu355 | -0.45 | Asp74 | -0.20 |
| 10 | Asn357 | -0.41 | Asp99 | -0.20 |

Top 10 ranked residues that involved in the binding of SARS-CoV-RBD and KIM1 Ig V domain were listed.

**Table S3.** Top 20 clinically identified mutations in SARS-CoV-2 spike protein.

| **Rank** | **Mutation** | **Occurrence (Log 10)** | **Rank** | **Mutation** | **Occurrence (Log 10)** |
|---|---|---|---|---|---|
| 1 | D614G | 4.786 | 11 | L54F | 2.575 |
| 2 | T723T | 3.507 | 12 | D839Y | 2.550 |
| 3 | Y789Y | 3.105 | 13 | G1044G | 2.549 |
| 4 | D936Y | 2.885 | 14 | T1100T | 2.513 |
| 5 | N824N | 2.806 | 15 | P715P | 2.473 |
| 6 | L5F | 2.797 | 16 | S477N | 2.453 |
| 7 | P1263L | 2.780 | 17 | V1008V | 2.343 |
| 8 | R21I | 2.738 | 18 | F541F | 2.337 |
| 9 | N439K | 2.730 | 19 | Q613Q | 2.303 |
| 10 | D294D | 2.714 | 20 | A879S | 2.303 |

Synonymous mutations are underlined. Data were acquired on August 25, 2020 from http://giorgilab.dyndns.org/coronapp/.

**Table S4.** Primers used in the study.

| Primers Name | Sequence: 5' to 3' |
|---|---|
| **Primers for qPCR** | |
| M-*Ace2*-F | CAAGTGTTGGCTTCGGTGTG |
| M-*Ace2*-R | ATTCAAGTGACCAGCGAGCA |
| M-*Kim1*-F | TCCACACATGTACCAACATCAA |
| M-*Kim1*-R | GTCACAGTGCCATTCCAGTC |
| M-*Actinb*-F | GGCTGTATTCCCCTCCATCG |
| M-*Actinb*-R | CCAGTTGGTAACAATGCCATGT |
| **Primers for CRISPR-Cas9 system** | |
| H-*KIM1*-KO-1-F | CACCGCTGACGGCCAATACCACTAA |
| H-*KIM1*-KO-1-R | AAACTTAGTGGTATTGGCCGTCAGC |
| H-*KIM1*-KO-2-F | CACCGGTTCGAACAGTCGTGACGGT |
| H-*KIM1*-KO-2-R | AAACACCGTCACGACTGTTCGAACC |
| H-*KIM1*-KO-3-F | CACCGGTCGTTGGAACAGTCGTCAT |
| H-*KIM1*-KO-3-R | AAACATGACGACTGTTCCAACGACC |

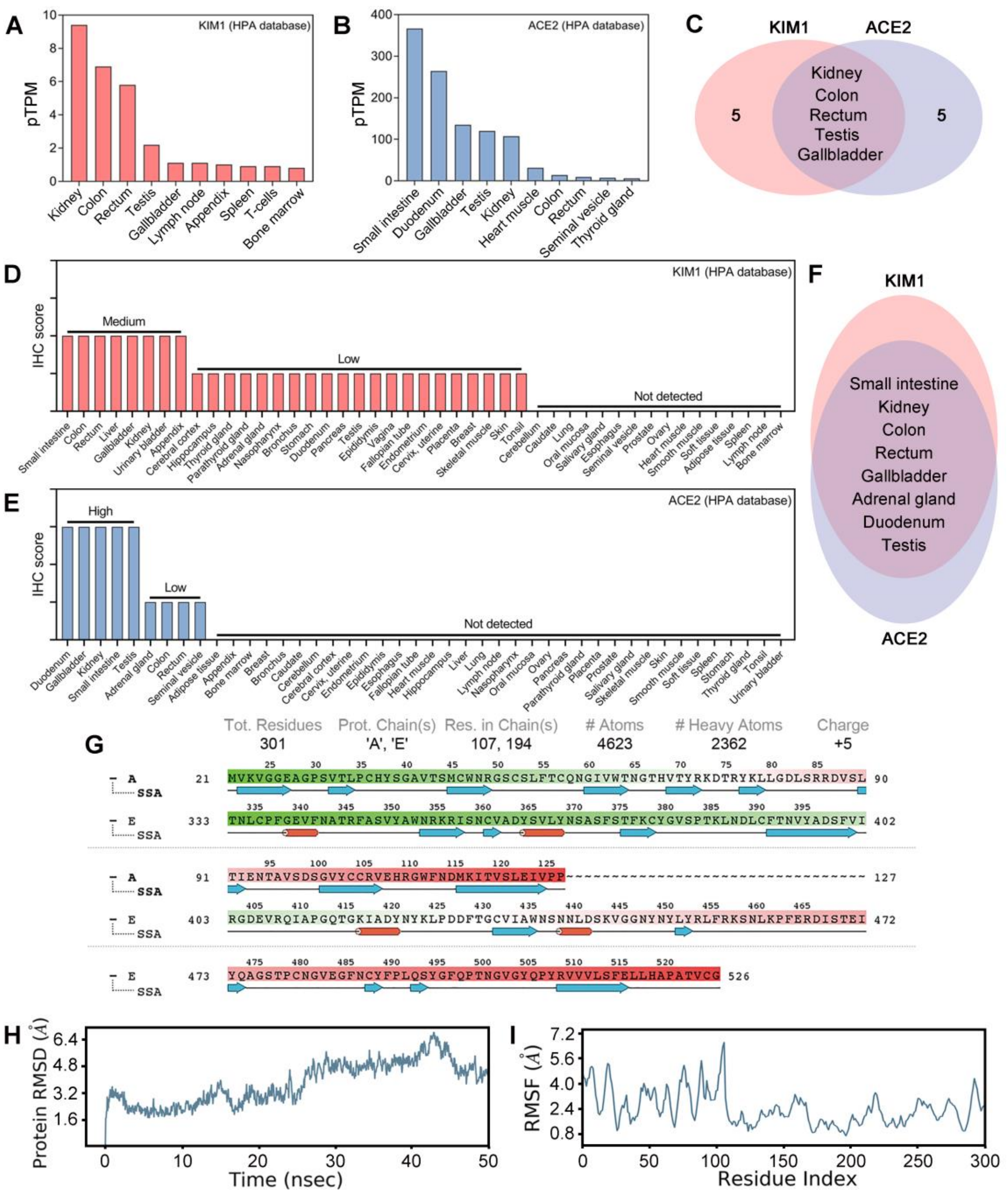


**Figure S1. Expression profiles of KIM1, ACE2 and molecular dynamics docking information of SARS-CoV-2-RBD and KIM1.** (A) Tissue transcriptional expression of *KIM1* from The Human Protein ATLAS dataset (HPA). pTPM, protein-coding transcripts per million. (B) Tissue transcriptional expression of *ACE2* from HPA. (C) Transcriptional expression intersection of *KIM1* and *ACE2* form HPA. (D) Histology-based protein expression levels of KIM1. (E) Histology-based protein expression levels of ACE2. (F) Histology-based protein expression intersection of KIM1 and ACE2. (G) Basic protein information of SARS-CoV-2-RBD and KIM1 Ig

V domain. Red cylinder represents α-Helix structure. Blue arrow indicates β-Sheet structure. (H) RMSD values of the SARS-CoV-2-RBD and KIM1 Ig V domain binding models. (I) RMSF values of the SARS-CoV-2-RBD and KIM1 Ig V domain binding models.

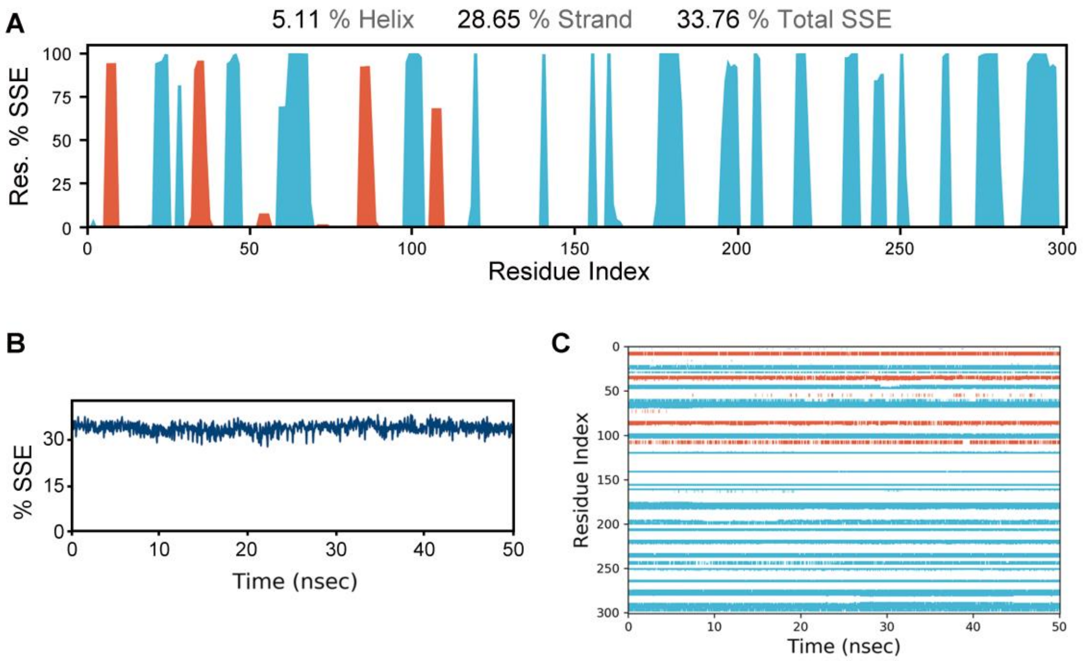


**Figure S2. Molecular dynamics simulations information of SARS-CoV-2-RBD and KIM1.** (A) Residue SSE distribution by residue index throughout the protein structure. Red column suggests α-Helix structure and blue colunm indicates β-strand structure. (B) SSE composition for trajectory frames in the dynamics simulation process of SARS-CoV-2-RBD and KIM1 Ig V domain, the curve indicates SSE composition for each trajectory frame over the course of the simulation. (C) SSE assignment of the residues over the simulation time, red dots indicates α-Helix structures and blue dots indicate β-strand structures. SSE, sencondary structure elements.

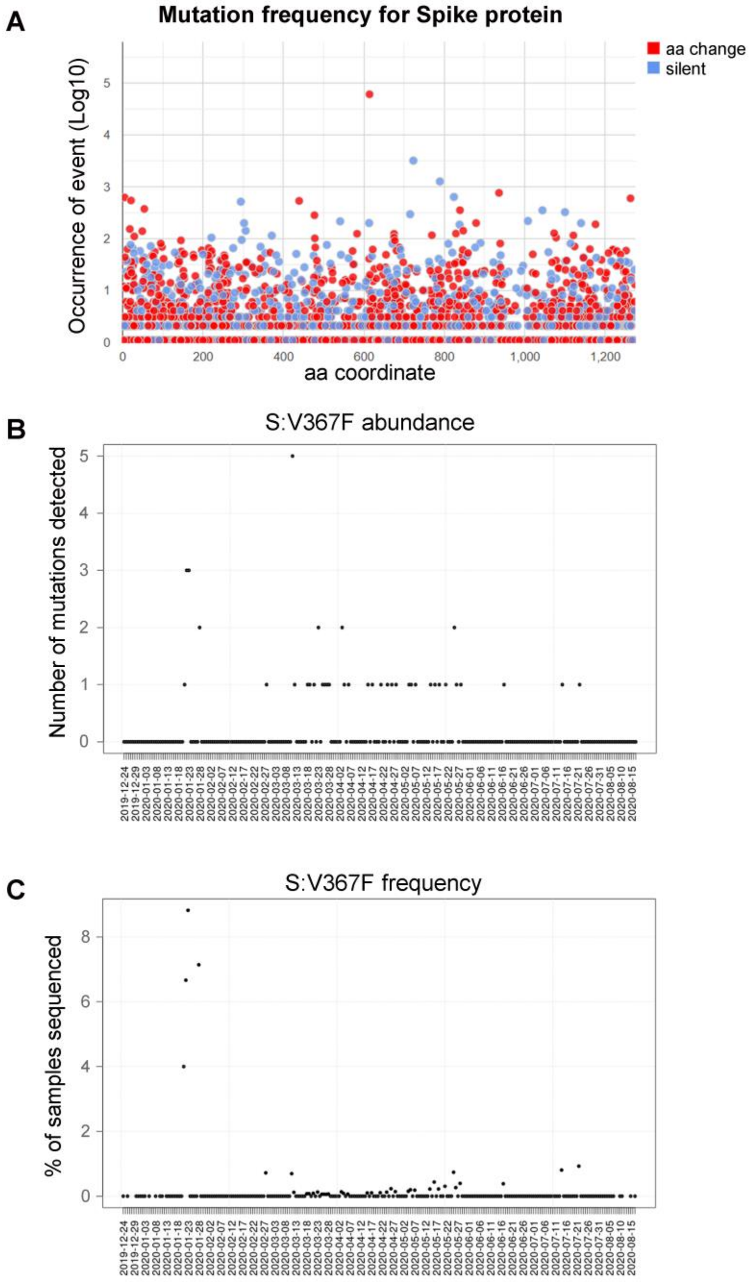


**Figure S3. Clinically identified mutations in SARS-CoV-2 spike protein**. (A) Mutation frequency for SARS-CoV-2 spike protein worldwide. Blue dots indicate synonymous mutation and red dots refer to amino acid substitution. (B) The number of clinically detected V367F mutation in COVID-19 patients. (C) Percentage of COVID-19 cases carrying V367F mutation. Data were acquired on August 25, 2020 from http://giorgilab.dyndns.org/coronapp/.

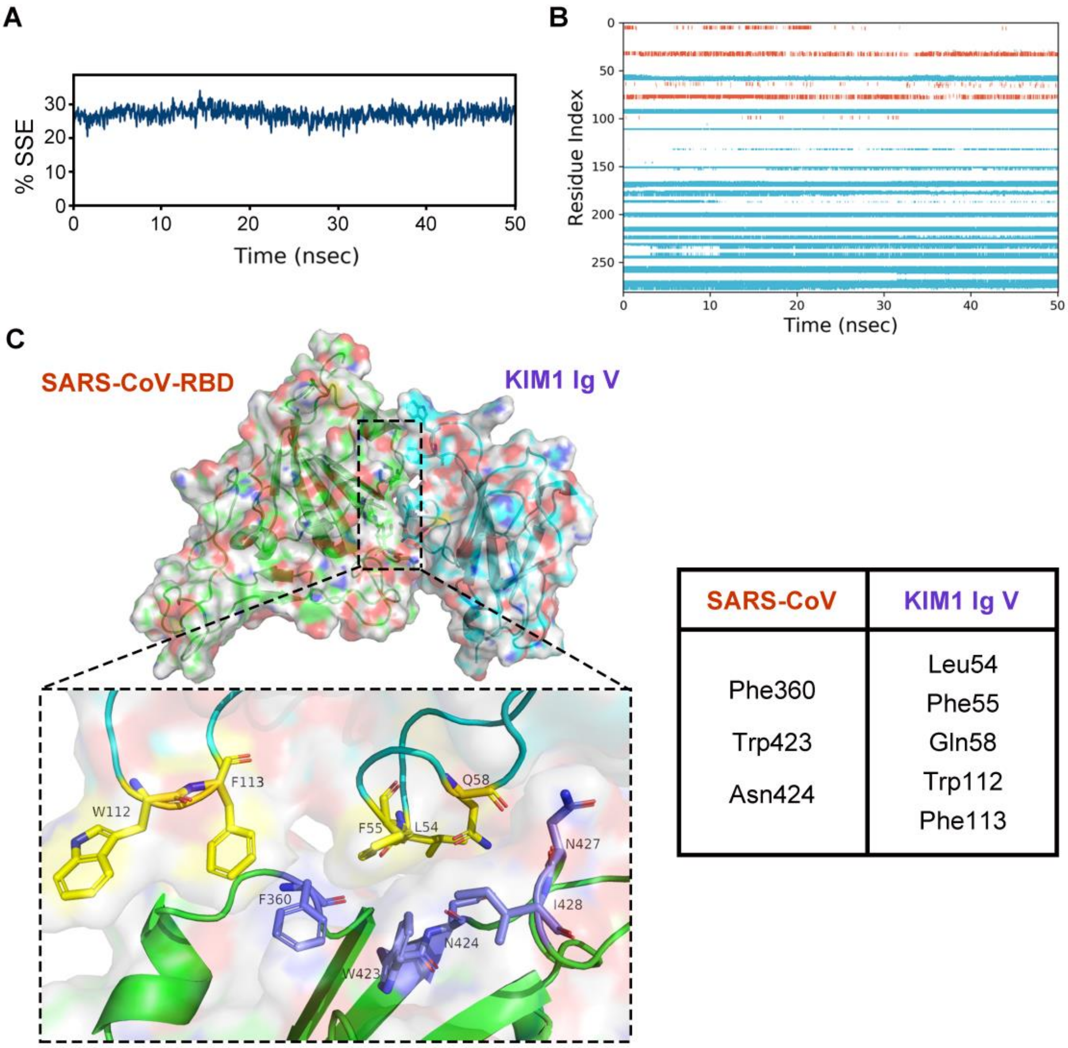


| SARS-CoV | KIM1 Ig V |
|---|---|
| Phe360<br>Trp423<br>Asn424 | Leu54<br>Phe55<br>Gln58<br>Trp112<br>Phe113 |

**Figure S4**. **Molecular dynamics simulations information of SARS-CoV-RBD and KIM1.** (A) *KIM1* mRNA expression in 10 SARS patients-derived peripheral blood mononuclear cells and 4 control samples (GSE1739). (B) Basic protein information of SARS-CoV-RBD and KIM1 Ig V domain. Red cylinder represents α-Helix structure of, blue arrow indicates β-Sheet structure. (C) RMSD values of the SARS-CoV-RBD and KIM1 Ig V domain binding models. (D) RMSF values of SARS-CoV-RBD and KIM1 Ig V domain. (E) Residue SSE distribution by residue index throughout the protein structure. Red column suggests α-Helix structure and blue colunm indicates β-strand structure. SSE, secondary structure elements.

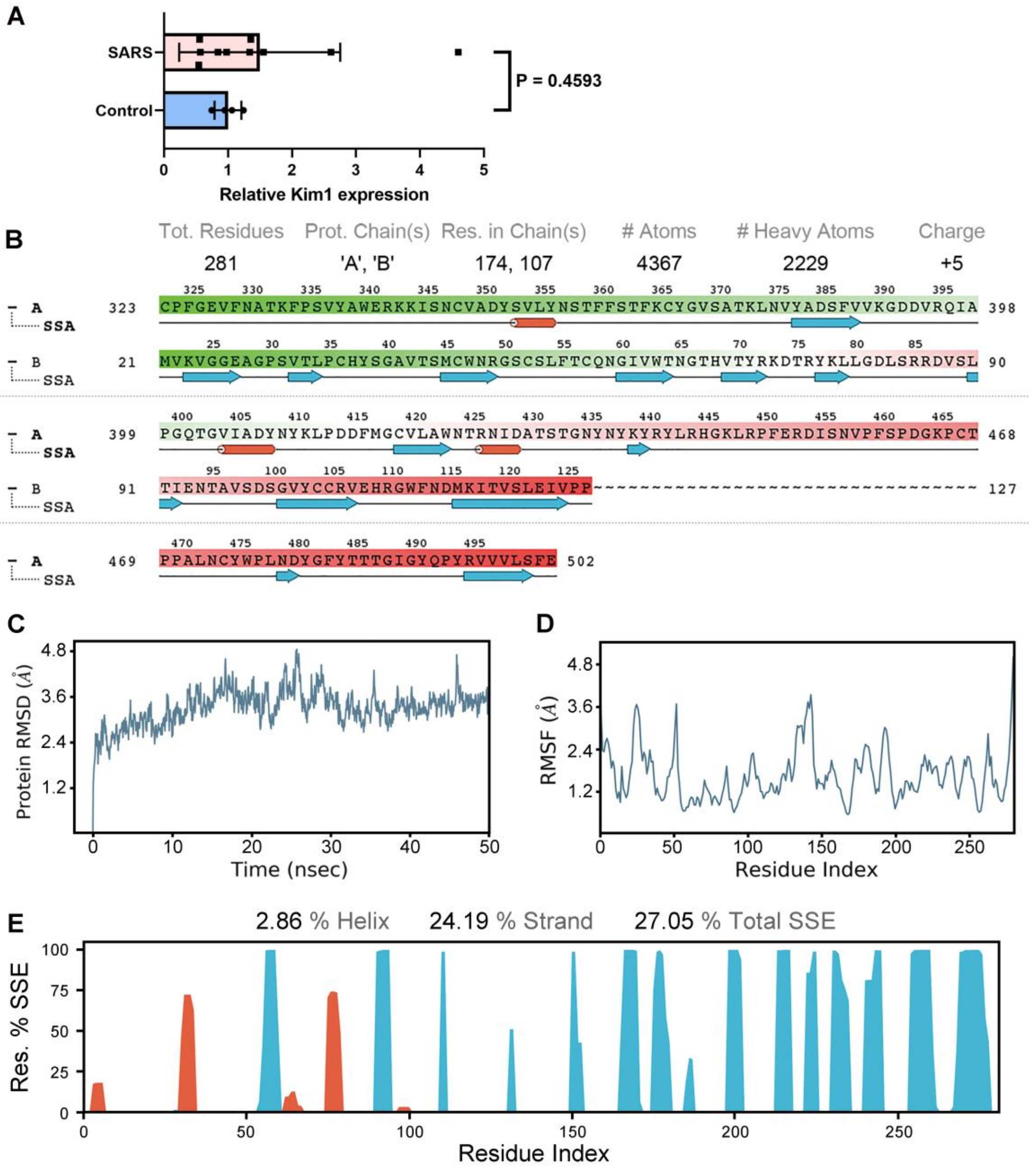


**Figure S5. Binding model of SARS-CoV-RBD and KIM1.** (A) SSE composition for trajectory frames in the dynamics simulation process of SARS-CoV-RBD and KIM1 Ig V domain, the curve indicates SSE composition for each trajectory frame over the course of the simulation. (B) SSE assignment of the residues over the simulation time, red dots indicate α-Helix structures and blue dots indicate β-strand structures. (C) Low-energy binding conformations of SARS-CoV-RBD binds to KIM1 Ig V domain. SSE, secondary structure elements.

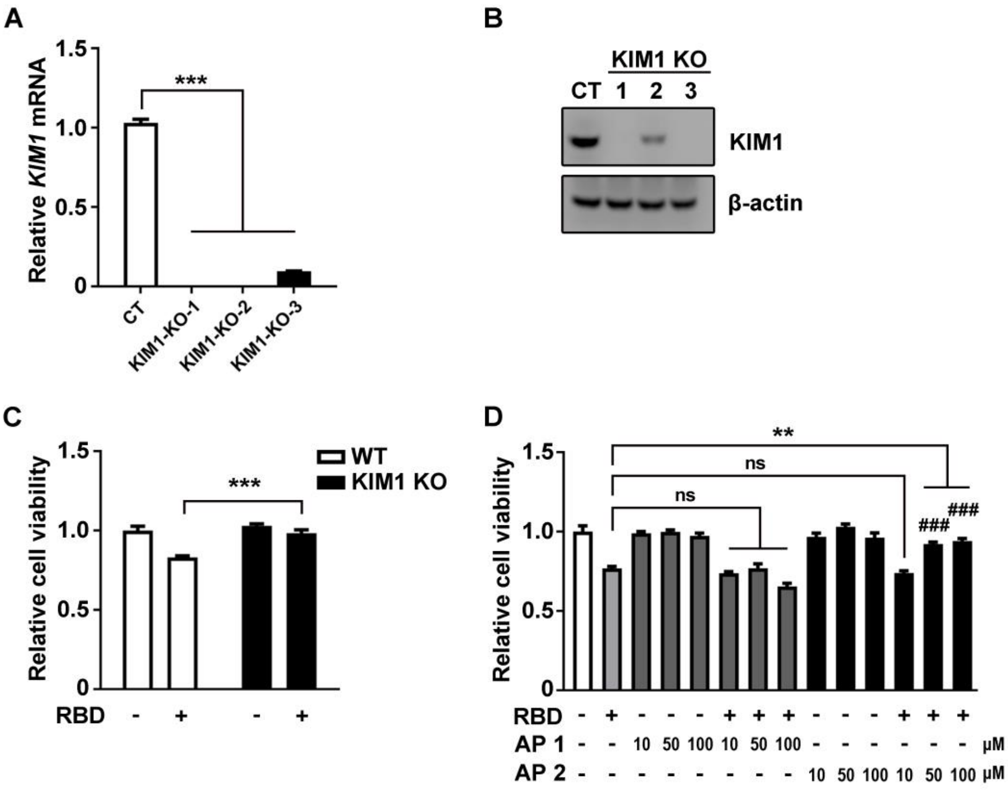


**Figure S6. Identification of KIM1 knockout HK-2 cell line and the protective effects of antagonist peptide on SARS-CoV-2-RBD induced cytotoxicity.** (A) *KIM1* mRNA expression in WT and KIM1 knockout HK-2 cell line. (B) KIM1 protein level in WT and KIM1 knockout HK-2 cell line. (C) Cytotoxicity of SARS-CoV-2-RBD in WT and KIM1 knockout HK-2 cell line. The concentration of SARS-CoV-2-RBD, 100 μg/ml. (D) Protective effects of antagonist peptide 2 against SARS-CoV-2-RBD. AP 1, antagonist peptide1. AP 2, antagonist peptide 2. $^{**}P < 0.01$, $^{***}P < 0.001$, $^{\#\#\#}P < 0.001$ compared to RBD + 10 μM AP2 group; ns, no significance.
